# Complex Hybridization in a Clade of Polytypic Salamanders (Plethodontidae: *Desmognathus*) Uncovered by Estimating Higher-Level Phylogenetic Networks

**DOI:** 10.1101/2024.01.29.577868

**Authors:** R. Alexander Pyron, Kyle A. O’Connell, Edward A. Myers, David A. Beamer, Hector Baños

**Affiliations:** Department of Biological Sciences, The George Washington University, Washington, DC 20052, USA; Department of Vertebrate Zoology, National Museum of Natural History, Smithsonian Institution, Washington, DC 20560-0162, USA; Deloitte Consulting LLP, Health Data and AI, Arlington, VA 22209, USA; Department of Herpetology, California Academy of Sciences, San Francisco, California 94118, USA; Office of Research, Economic Development and Engagement, East Carolina University, Greenville, NC 27858, USA; Department of Biochemistry and Molecular Biology, Faculty of Medicine, Dalhousie University, Halifax, NS B3H 4R2, CA; Department of Mathematics and Statistics, Faculty of Science, Dalhousie University, Halifax, NS B3H 4R2, CA; Department of Mathematics, California State University San Bernardino, San Bernardino, CA, USA

**Keywords:** Phylogenetic networks, Repeated hybridization, Ecological speciation, *Desmognathus* salamanders, Reticulate evolution

## Abstract

Hybridization between incipient lineages is a common feature of ecomorphological diversification. We examine these phenomena in the Pisgah clade of *Desmognathus* salamanders from the southern Appalachian Mountains of the eastern United States. The group contains four to seven species exhibiting two discrete phenotypes, aquatic “shovel-nosed” and semi-aquatic “black-bellied” forms. These ecomorphologies are ancient and have apparently been transmitted repeatedly between lineages through introgression. Geographically proximate populations of both phenotypes exhibit admixture, and at least two black-bellied lineages have been produced via reticulations between shovel-nosed parentals, suggesting complex transmission dynamics. However, computational constraints currently limit our ability to reconstruct network radiations from gene-tree data. Available methods are limited to level-1 networks wherein reticulations do not share edges, and higher-level networks may be non-identifiable in many cases. We present a heuristic approach to recover information from higher-level networks across a range of potentially identifiable empirical scenarios, supported by theory and simulation. When extrinsic information indicating the location and direction of hybridization events is available, our method can yield successful estimates of non-level-1 networks, or at least a reduced possible set thereof. Phylogenomic data strongly support a single backbone topology with up to five overlapping hybrid edges. These results suggest an unusual mechanism of ecomorphological hybrid speciation, wherein a binary threshold trait causes hybrids to shift between two microhabitat niches, promoting ecological divergence between sympatric hybrids and parentals. This contrasts with other well-known systems in which hybrids exhibit intermediate, novel, or transgressive phenotypes. Finally, the genetic basis of these phenotypes is unclear and further data are needed to clarify the evolutionary basis of morphological changes with ecological consequences.

## Introduction

Hybridization between lineages is increasingly recognized as a common feature of evolutionary history, now frequently represented as phylogenetic networks (Solís-Lemus and Ané 2016). Additionally, speciation is often marked by a complex interplay between historical isolation processes and genomic introgression (Nosil 2008; Abott et al. 2014). Consequently, intricate phylogenetic networks with multiple hybrid edges are likely to be a common feature of species radiations (e.g., Gante et al. 2016). In emerging lineages, species boundaries can remain porous for long periods of time (Harrison and Larson 2014), and multiple contacts between incipient species can lead to multiple instances of admixture (Barth et al. 2020). Repeated introgression can therefore blur species boundaries (Crow et al. 2007; Minder and Widmer 2008) and produce species histories not adequately represented by bifurcating trees (Huson and Bryant 2006; Solís-Lemus and Ané 2016). These repeated contacts may themselves drive evolutionary diversification through hybridization and genetic transfer (Seehausen 2004; Rosenblum et al. 2012). The transmission of adaptive alleles and quantitative trait loci via introgression may also induce ecological speciation as hybrids develop novel phenotypes, occupy new adaptive landscapes, or switch between ecomorphs (Hatfield and Schluter 1999; Nolte et al. 2005; Rougemont et al. 2015; Makowicz and Travis 2020; Short and Streisfeld 2023).

A key example are *Desmognathus* salamanders, which exhibit complex reticulation patterns that frustrate ordinary phylogenetic inference. Our recent studies (Pyron et al. 2020, 2022) revealed at least 47 candidate lineages in a genus previously containing only 21 described species. These taxa display extensive genealogical discordance and complex histories of hybridization. Remarkably, they are both polytypic in the sense that recent radiations can display multiple distinct phenotypes within and among species, and cryptic in the sense that distantly related lineages often share highly similar phenotypes (Kozak et al. 2005; Beamer and Lamb 2008). Of these, black-bellied (previously called *D. “quadramaculatus”*) and shovel-nosed (*D. marmoratus*) salamanders were long thought to represent only two independent species, but instead comprise 11 distinct lineages in two separate clades – 8 of which are now recognized as species – with repeated phylogenetic emergence of their two distinctive phenotypes (Kozak et al. 2005; Pyron et al. 2022). The Nantahala and Pisgah clades (see Jones and Weisrock 2018) both contain black-bellied and shovel-nosed species but are distantly related and were apparently involved in a deep-time reticulation that transferred mitochondrial haplotypes and potentially alleles for each phenotype (Jackson 2005; Pyron et al. 2020).

Here, we focus on the Pisgah clade of four black-bellied (heavy-bodied and semi-aquatic) and three shovel-nosed (slender and aquatic) lineages (Fig. 2, Fig. S1–3) from the southern Appalachian Mountains of the eastern United States. The group shared a common ancestor ∼3 Ma in the southern Appalachian Mountains (Kozak et al. 2005). Four of the seven lineages represent recognized species: the black-bellied *Desmognathus kanawha* and *D. mavrokoilius,* and the shovel-nosed *D. intermedius* and *D. marmoratus.* The other three are geographic genetic lineages without names, the shovel-nosed *“marm”* G, and the black-bellied *“quad”* C and *“quad”* G, all named as such for historical reasons (see Supplemental Information at Zenodo repository https://doi.org/10.5281/zenodo.10386850), with apparent hybrid ancestry from multiple parental lineages (Pyron and Beamer 2022; Pyron et al. 2022). These seven taxa show variable relationships to each other using mitochondrial and nuclear datasets, and recent analyses suggest extensive hybridization – potentially both ancestral and ongoing – between numerous populations and lineages in the group. We hypothesize that several of the genetically and morphologically distinct populations represent ecomorphologically mediated “hybrid species” from ancestrally differentiated parental lineages (see Schumer et al. 2014).

By this, we follow Mallet (2007) in meaning “cases where hybrid allelic combinations contribute to the spread and maintenance of stabilized hybrid lineages generally recognized as species.” Furthermore, if we “recognize species as multi-locus ‘genotypic clusters’ … a hybrid species will then be a third cluster of genotypes, a hybrid form that has become stabilized and remains distinct when in contact with either parent.” A key feature for success in such instances is thought to be ecological differentiation from parental lineages (Abbott et al. 2010), typically through intermediate, novel, or transgressive phenotypes. In simple instances with two parentals and one hybrid, this may be easy to demonstrate (e.g., Gompert et al. 2006). Alternatively, the repeated evolution or vertical transmission from standing ancestral variation of alternative ecomorphs may facilitate ecological segregation of parentals and hybrids over repeated instances of introgression (De-Kayne et al. 2022; Short and Streisfeld 2023). One might also draw a distinction between ancient, well-established or ‘stabilized’ hybrid lineages or species and those arising from more recent secondary contact and subsequent admixture (Chaturvedi et al. 2020). In more complex scenarios with multiple reticulations, however, current limitations on tree-based network analyses limit our ability to estimate these historical processes.

We use SNP data from genotype-by-sequencing (GBS, Elshire et al. 2011) and gene trees from anchored hybrid enrichment (AHE; Lemmon et al. 2012) loci to interrogate the phylogeny of the Pisgah clade, with a geographically well-sampled and morphologically vouchered dataset to answer several questions. First, what network best describes relationships among the seven species-level lineages? Second, which present-day populations represent parentals, introgressed descendants, or hybrid species? Third, do patterns of admixture and network relationships indicate a role for a phenotypically dimorphic threshold trait promoting hybrid ecomorphological speciation? To do so, we propose a new heuristic pipeline based on coalescent theory and supported by simulations that can recover signal from higher-level networks with multiple intersecting cycles from gene-tree data (at least in part) when combined with other indicators of the location of these edges, while accounting for incomplete lineage sorting under the network multispecies coalescent (NMSC; Meng and Kubatko 2009; Yu et al. 2011; Zhu et al. 2016).

The NMSC model describes the formation of gene trees within networks in the presence of both ILS and hybridization or other discrete horizontal gene transfer events. There are many useful methods to detect hybridization under the NMSC. Current inference methods include pseudo-likelihood, Bayesian, and combinatorial methods (Steel 2016; Degnan 2018; Hahn and Hibbins 2019). For the first class, there are no identifiable results beyond level-1 networks; for the second class, only small datasets have been analyzed due to their high computational cost; and for the third class, there are many estimates of higher-level networks that do not consider ILS or are restricted to level-1 if they do. Given the complexity of the NMSC, statistical identifiability of networks has been achieved for only a small class of networks known as “level-1” (Solís-Lemus and Ané 2016; Allman et al. 2019; Baños 2019). A binary network is defined as level-1 if no two cycles (loops of branches formed by a reticulation) share an edge (Rosselló and Valiente 2009). This excludes networks that contain hybrids arising directly from other hybrids, or networks containing one ancestor that is directly involved in at least two different hybridization events, a potentially common empirical result for network radiations of rapidly radiating species (Gante et al. 2016; Kozak et al. 2021).

Based on recent results on identifiability under the NSMC (Allman et al. 2023), we leverage asymmetries in the empirical frequency of concordance factors estimated from gene trees to isolate a set of candidate backbone trees for the network that are congruent with the dominant (most frequent) quartets. From these, we select one (or a consensus) as a backbone using heuristic information from previous species-tree analyses. To this backbone, we add hybrid edges using additional heuristic information regarding the likely topological location and direction of gene flow, such as genomic tests of excess allele-sharing and mitochondrial introgression. This yields a complex (non-level-1) phylogenetic network that is mathematically congruent with most of the empirical gene trees and qualitatively concordant with additional heuristic sources of information regarding the most likely placement and direction of hybrid edges (Fig. 1). We provide preliminary support and validation for this approach via two simulations: one of the simplest-possible case and one resembling our empirical scenario.

**Figure 1.**
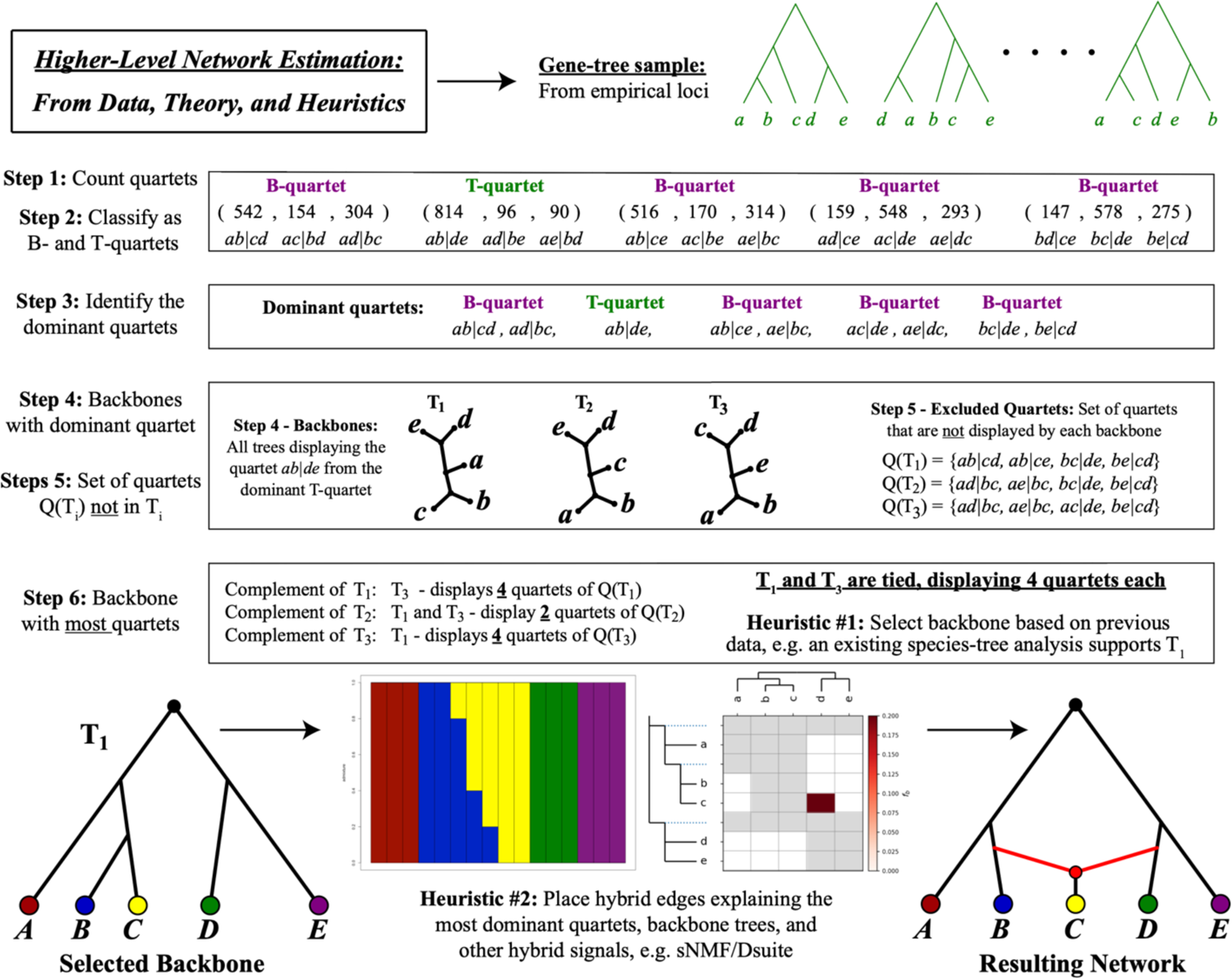
An overview schematic of the steps taken in our heuristic approach to estimating potential resolutions for higher-level networks. From a sample of gene trees, complete the first 2 steps outlined above using the NANUQ function from MSCquartets (Rhodes et al. 2021). The remaining steps are performed using custom code available in the SI and associated repositories (see Data Availability Statement) to arrive at a set of candidate backbones that are congruent with the dominant quartets. Using heuristic information, one (or a consensus) tree is selected as a backbone, to which hybrid edges are then added using additional heuristic information such as individual ancestry coefficients or excess allele-sharing. These edges are added manually in such a way as to reconcile each other and the candidate backbones while explaining the largest number of dominant quartets (see SI), yielding a non-level-1 network that satisfies the greatest amount of constraint and hybrid signal reconciled from various empirical sources.

**Figure 2.**
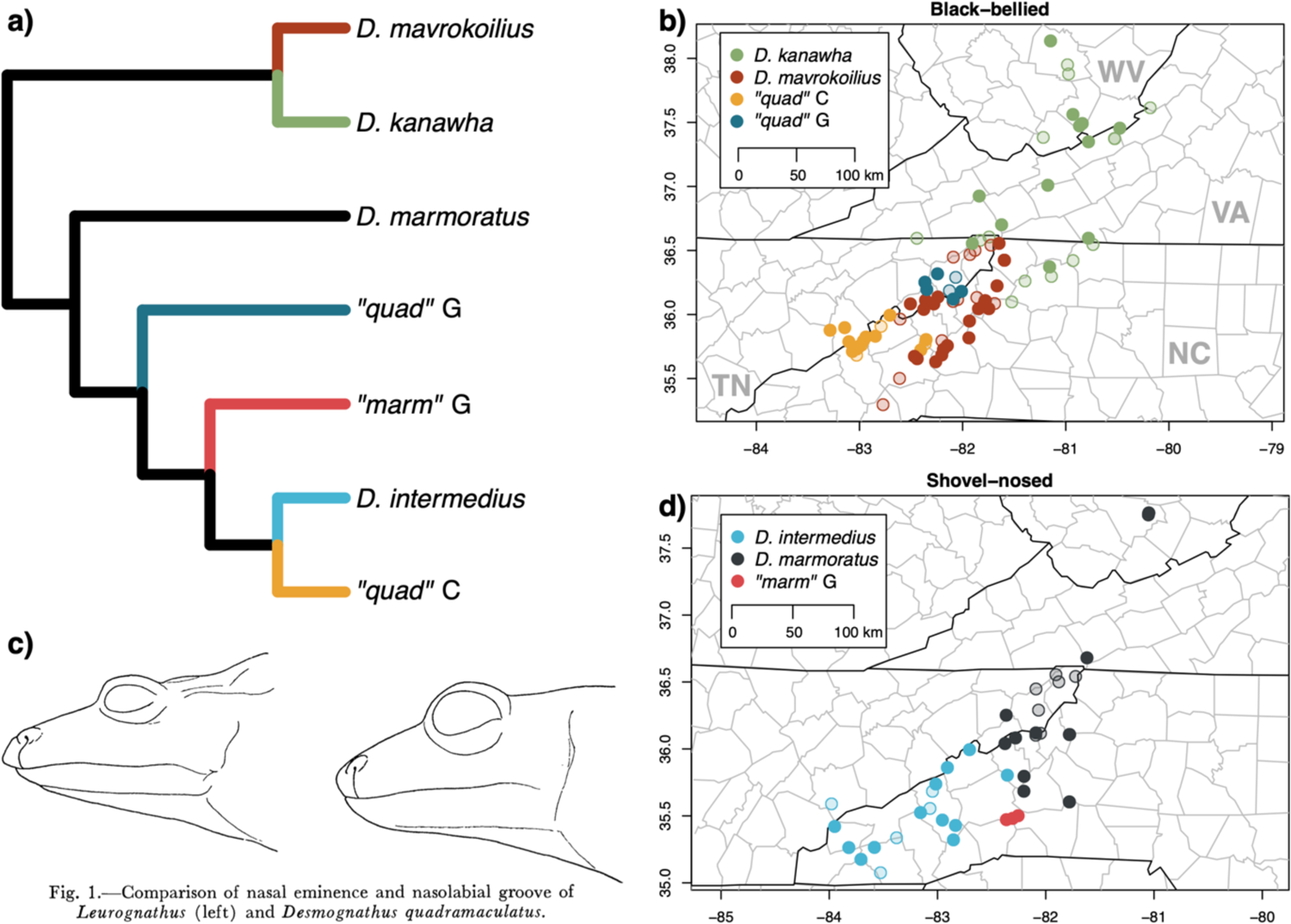
Tree-based estimates of the phylogenetic relationships between the seven Pisgah-clade lineages (a; Pyron et al. 2022), maps (b, d) of the 335 specimens from 105 localities grouped by phenotype (b: shovel-nosed, d: black-bellied; Appendix S2) and lineage, and a historical illustration (c) of the binary phenotype as expressed in head shape and snout profile (from Martof 1962). State abbreviations in (b) are WV – West Virginia, VA – Virginia, NC – North Carolina, and TN –Tennessee. Solid dots indicate GBS data (69 sites, 77 individuals), while semi-transparent localities were sampled for morphology only (36 sites).

We find congruent support for a higher-level network that is supported by admixture patterns from SNP-based methods and mitogenomic capture. This topology suggests that 3–5 of the seven lineages are of hybrid origin between genetically and phenotypically differentiated ancestral parental populations. The binary phenotype is seemingly polygenic but is not intermediate in admixed individuals. This contrasts with the more common pattern in which hybrids are intermediate, novel, or transgressive with respect to parentals. We hypothesize that the ecological separation of these phenotypes is responsible for the maintenance of newly arising hybrid lineages, an unusual mechanism of ecomorphological sympatric hybrid speciation.

Finally, both phenotypes – which may comprise subtly distinct morphologies in each lineage – were apparently captured during ancient introgression from the distantly related Nantahala clade, a pattern reflected in the mitochondria but not yet recovered in the nuclear genome (Pyron et al. 2020, 2022). Use of our new pipeline at deeper scales may allow such patterns to be recovered in future analyses. We tentatively allocate *“marm”* G to *D. marmoratus,* but the evolutionary origins and taxonomic status of *“quad”* C and G remain to be unraveled further.

## Materials and Methods

### Higher-Level Network Inference

First, we provide a detailed explanation of a new heuristic approach for obtaining structural information for networks from quartet data derived from gene trees (Fig. 1). This information is then combined with external evidence of hybridization to estimate a network heuristically. In our empirical case, we rely on i) previous evidence of mitochondrial haplotype interchange between named lineages (Kozak et al. 2005; Beamer and Lamb 2020; Pyron et al. 2020), ii) individual ancestry coefficients from sNMF analyses (Frichot et al. 2014) suggesting admixture, and iii) excess allele-sharing under the MSC model from a Dsuite analysis (Malinsky et al. 2021). While the latter two might be susceptible to confounding evolutionary and population-genetic processes (e.g., rate variation) and decreased power with multiple reticulations (e.g., Frankel and Ané 2023), they offer at least preliminary external heuristics.

First, we describe the procedure, then provide the empirical results from the AHE and GBS datasets. Additional detail is given in the SI (see Tables S1–2). The basis of our method relies on *Theorem 1* (see Allman et al. 2023), which is stated below. We say that a network *N* displays a tree *T* if *T* can be obtained from *N* after removing some hybrid edges. A *quartet network Q* of *N* is a subnetwork of *N* on 4 taxa. Following Allman et al. (2023), we say that an unrooted quartet *Q* on network X is a B-quartet network if there is no edge in *Q* that induces a split of the form *xy*|*zw*, for taxa *x, y, z, w* in *X;* otherwise, we say *Q* is a T-quartet. Roughly speaking, a B-quartet network is a network that contains a hybridization involving the 4 taxa together; one might think of these as “deep-hybrids” quartets. It can be shown that a T-quartet network (or “non-deep-hybrid”) displays only one tree topology, while a B-quartet network displays at least two. Figure S10 depicts both a B-quartet network and a T-quartet network. The network on the left of Figure S9 contains the set of all unrooted trees displayed in Figure S11.

**Theorem 1:** Let N be a rooted species phylogenetic network. Then under the NMSC for generic parameters, the collection of unrooted B-quartet networks can be identified by the ordering of concordance factors. Specifically, a quartet network is a B-quartet if and only if the triple of concordance factors does not contain two equal entries (see Allman et al. 2023).

It was previously shown that for level-1 networks one can determine the collection of B-quartets (Solís-Lemus et al. 2017; Baños 2019). Theorem 1 is a generalization of this result for arbitrary networks. Nonetheless, for an arbitrary network the structural properties of the B-quartet subnetworks are not necessarily identifiable from only concordance factor orderings, as in the level-1 case. Let *N* be a rooted network on taxon set *X* and let *a, b, c,* and *d* be four distinct taxa in *X*. Let *A, B, C,* and *D* be genes sampled from species *a, b, c,* and *d,* respectively. Given a gene quartet *AB|CD*, the quartet concordance factor *CF_AB|CD_* is the probability under the NMSC on *N* that a gene tree displays the quartet *AB|CD*, and *CF_abcd_* = (*CF_AB|CD_, CF_AC|BD_, CF_AD|BC_*) is the ordered triple of concordance factors (CFs) of each quartet on the taxa *a, b, c, d*. Our method uses the information from CFs to recover displayed trees of a network.

A key component of our heuristic method is to associate quartet trees with their induced quartet networks. Such quartet trees are determined by the triplet of concordance factors. As mentioned, any B-quartet network *Q* displays at least two of the three 4-taxon unrooted trees. We assume that these are the trees associated with the two largest concordance factors. That is, we assume that the topologies of the two most probable gene-tree quartets agree with the topology of the species network. With each B-quartet network *Q*, we associate two quartet trees: those defined by the two largest CFs. In a similar fashion, each T-quartet network *Q* can be associated with a single quartet tree, the one corresponding to the concordance factor that is unequal to the other two in the triplet of concordance factors (two entries of the triplet CF must be equal from Theorem 1). It can be shown that this tree is the one defined by the non-trivial split induced from the definition of a B-quartet. Moreover, it can be shown that this tree topology is the only quartet tree displayed by the quartet network.

Based on *Theorem 1,* which states that by looking only at gene-tree topology frequencies on a 4-taxon network, we can determine if the species network generically displays only one topology (a T-quartet) and which one, or if it displays multiple tree topologies (B-quartets) and which are likely the two most dominant. For the heuristic portion, we construct sets of trees that display such collections of quartets. Such trees are likely to be features of the network; in most cases, species networks will display the topology of the most dominant gene tree quartets. We conduct two simulation studies demonstrating the effectiveness of our method, described below and in the SI. Our heuristic approach to estimating non-level-1 networks proceeds as follows:

1. For every subset of 4 taxa, determine the empirical quartet counts (empirical concordance factors) from the gene trees.
2. Apply a statistical hypothesis test to the empirical concordance factors as in Mitchell et al. (2019) to determine which quartet networks are B-quartet networks and which are T-quartet networks.
3. As detailed in the SI, associate one quartet tree with each quartet identified as a T-quartet, and associate two quartet trees with each quartet identified as a B-quartet. We expect that the collection of all quartet trees associated with B-quartets and T-quartets will be displayed in the network. We refer to this collection as the set of dominant quartet trees.
4. Construct a set *T^0^* consisting of all trees that display only dominant quartet trees (those quartet trees in step 3). Any tree in *T^0^* could potentially be displayed in the network since, for any subset of 4 taxa, each of these trees will display a dominant quartet tree on those 4 taxa. This step is done exhaustively by looking at all possible trees displaying only dominant quartets.
5. For each tree *T* in *T^0^*construct a set *Q(T)* consisting of the set of quartet trees associated with all dominant quartets B-quartets which are not displayed in *T*. Note that for any candidate tree *T* in *T^0^*, *Q(T)* contains all the dominant quartet trees that are not displayed in *T*. If one would assume that *T* is a displayed tree in the network, then there should be other displayed trees in the network that display the remaining dominant quartet trees, i.e. those in *Q(T)*. Loosely speaking, *Q(T)* is the information in the network not accounted for by T.
6. For each tree *T* in *T^0^*, find the set of trees *S(T)* contained in *T^0^* that display the maximum number of quartet trees in *Q(T)*. The output of the method consists of those trees in *T^0^* such that the trees in *S(T)* displayed the greatest number of quartets for all *T* in *T^0^*, and their associated trees are the set *S(T)*. We refer to *S(T)* as the complement of *T*. In particular, note that for any candidate tree *T* in *T^0^*, the set *S(T)* contains the trees that display most of the dominant quartet trees not displayed by *T*. One can think of *S(T)* as the set of candidate trees that display most of the information missing in *T*. The output consists in those trees *T* whose complement describes most of the missing data in *T*, together with *S(T)*.

From the output of the function ‘NANUQ’ in the *R* package MSCquartets (Rhodes et al. 2021) one can obtain: (a) the set of quartet trees associated with quartets identified as T-quartet networks, and (b) the two quartet trees associated with the two largest concordance factors of *Q*, for all quartets *Q* identified as a B-quartet network. Steps 3–6 can be implemented in many ways using distinct combinatorial results (e.g., Bryant and Steel 2001; Semple and Steel 2003; Reaz et al. 2014; Rhodes 2020). Since the dataset of interest here has only seven taxa, these steps can be done quickly by doing an exhaustive search on all 945 unrooted seven-taxon tree topologies. We provide a simulation and demonstration of the method to showcase its empirical efficacy. This paves the way for new theoretical results on mathematical identifiability, hopefully to be followed by new methods to construct higher-level networks within the adequate family.

We conducted two simulations to test this approach. First, we show each step using a simulated data set from a level-1 network of 5 taxa and one hybridization event (Fig. 1; SI). We simulated 1000 gene trees from the network using Hybrid-Lambda (Zhu et al. 2015). Note that since this is a 5-taxon network, there are 5 possible quartets, and there are 15 possible unrooted trees displaying all 5 taxa. Following the six steps yields a strongly supported backbone estimated from the signal in the gene trees alone. Adding hypothetical heuristic information showing admixture between *B* and *C* and excess allele-sharing between *C* and *D* allows for placement of the two hybrid edges yielding the “correct” network.

We then proceeded with a complex example approximating our empirical dimensions. The simulated data *G* consists of 5000 gene trees generated using Hybrid Lambda (Zhu et al. 2015) using the level-2 network *N* depicted on the left of Figure S9. Note that since this network contains 7 taxa, it contains _7_C_4_ = 35 quartet networks. Also, recall that there are (7 × 2-5)!! = 945 distinct unrooted binary tree topologies relating to 7 taxa (Figs. S9–11). For *G*, NANUQ shows that an assumption is being violated, since the network is not level-1. We find that 8 of the 35 quartets were identified as T-quartets and 27 quartets were identified as B-quartets. For step 3, we used the output from the R function nanuq. The quartet trees associated with T-quartets are shown in Table S1, while the quartet trees associated with B-quartets are shown in Table S2. For step 4, we found that there are 45 trees displaying all 8 quartet trees in Table S1. Of the 45 trees found, only 13 agree with the B-quartet signals. That is, only 13 trees display exactly one quartet tree per row of Table S2. Therefore, *T^0^*consists of 13 such trees. For steps 5 and 6, there are 4 out of the 13 trees in *T^0^* whose trees in *S(T)* display 24 quartets, the maximum number of quartets displayed by any tree in *S(T)* for any tree in *T^0^*. These 4 trees are depicted in Figure S11. For the other 9 trees, their corresponding trees in *S(T)* display between 20 and 22 quartets. Note that the trees in Figure S11 are all possible unrooted trees obtained from *N* by removing one hybrid edge per hybridization event. Let *t_1_*, *t_2_*, *t_3_*, and *t_4_* be such trees in order of appearance from left to right. Specifically, we obtained *S(t_1_) = {t_4_}, S(t_2_) = {t_3_}, S(t_3_) = {t_2_}, S(t_4_) = {t_1_}*. Therefore, no more trees need to be considered and the output is exactly those 4 trees. The output obtained from this example is enough to reconstruct the unrooted version of *N*.

This methodological approach appears to yield meaningful signals of networks beyond level-1 under the NMSC, and in some cases these signals are enough to recover the full network. Theoretically, our method is consistent with level-1 networks except in very rare cases related to 3_2-cycles as detailed in Allman et al. (2019). We expect these cases to remain problematic in more general networks. Moreover, other problematic scenarios may arise from more complex hybridization events. One of the main limitations in determining to what extent these problems persist and appear is the lack of identifiability results and study of complex networks. We hypothesize that in biological settings, these events correspond to unrealistically extreme parameters as in the level-1 case (for example having ‘massive population sizes’ or ‘close to zero number of generations’). Therefore, our method should perform adequately in many cases.

### Salamander Networks

To address the complex hybrid origin of the seven lineages in the Pisgah clade of *Desmognathus*, we applied our new method to an existing densely sampled dataset comprising 233 gene trees and 122 individuals from 108 populations (Pyron et al. 2022). These gene trees were estimated from long (1,025–5,24bp) Anchored Hybrid Enrichment loci (AHE; Lemmon et al. 2012), which are designed to species-level variation and yield well-resolved topologies. We began with the AHE dataset by condensing the topological arrangement of individuals down to species-level representation by doing the following:

1. We selected one individual at random for each species.
2. We kept a copy of each gene tree that displays these individuals and removed all other individuals. This left a set of 233 gene trees with seven taxa each.
3. We repeated this procedure 10,000 times resulting in 1,944,981 gene trees. This procedure consolidated signals found in different individuals (e.g. different levels of introgression across geographic space) and reduced variability between species. Many gene trees exhibit redundancy, but we do not expect this to bias results because we are estimating concordance factor rankings.
4. We then ran NANUQ on this set of gene trees and observed, as expected, a violation of the hypotheses. Therefore, we performed our method as outlined above under Steps 1–6.

Finally, we combined the signals from the pipeline’s output trees (i.e., the candidate backbones), Dsuite, sNMF, and the estimated B-quartets to infer a phylogenetic network. First, we fixed the backbone tree (Figure S12, left) obtained from our method in Step 6, which is identical to the topology previously estimated for the concatenated and species-tree analysis of these AHE data (Pyron et al. 2020, 2022) and very similar to the mitochondrial estimate (Beamer and Lamb 2020). We then added hybrid edges heuristically in the order of the most congruent signals (i.e., encompassing the greatest number of quartets) for observed B-quartets vs. T-quartets. We based these edges, and when possible, their orientation, on additional sources of information we considered credible to indicate the existence of introgression events (see below), together with the information obtained from B-quartets to find the signals agreeing with the greatest number of B-quartets estimated in Step 2. Consequently, we choose hybrid edges and directions for which the resulting network displays the greatest number of the quartet trees associated with B-quartets and align with the external data.

These data were obtained from three sources. The first is mitochondrial exchange indicated in previous datasets (Kozak et al. 2005; Beamer and Lamb 2020; Pyron et al. 2020). The second are admixture estimates from our previous analyses of the AHE data (Pyron et al. 2022) using sNMF (Frichot et al. 2014) and Dsuite (Malinsky et al. 2021) with a threshold of 20% shared ancestry interpreted as significant. Third, to augment these analyses with a broader genome-wide scan of SNPs, increase targeted sampling of populations, and avoid circularity, we generated an additional dataset for these lineages using genotype-by-sequencing (GBS; Elshire et al. 2011) analyzed using Dsuite and sNMF. The potential hybrid events indicated by these analyses are summarized in Table S5. A minor signal of excess allele-sharing between *D. kanawha* and *D. marmoratus* in the Dsuite analysis is likely artifactual, resulting from the stronger and more likely *D. marmoratus/D. mavrokoilius* hybrid edge (see Malinsky et al. 2021; Pyron et al. 2022). Gene flow between *D. intermedius* and *“quad”* C, and between *D. kanawha* and *D. mavrokoilius* was previously indicated by mitochondrial exchange (Beamer and Lamb 2020) and admixture analysis (Pyron et al. 2022). However, as both pairs are sister lineages and may represent the signal of primary divergence, this consequently does not affect the network.

### Sampling and Sequencing

We sampled 77 individuals from the seven lineages (*n* = 3–21) in the Pisgah clade (Appendix S1): the black-bellied *Desmognathus kanawha* (*n* = 12), *D. mavrokoilius* (*n* = 21), and the enigmatic *“quad”* C (*n* = 12) and *“quad”* G (*n* = 6) lineages of putative hybrid origin; and the shovel-nosed *D. intermedius* (*n* = 11)*, D. marmoratus* (*n* = 12), and the *“marm”* G (*n* = 3) lineage with complex ancestry (Fig. S1–3). These were each reciprocally monophyletic and phenotypically monomorphic in our previous phylogenetic analyses (Pyron et al. 2020), but patterns of genomic composition varied between the *“quad”* C and G and *“marm”* G lineages in admixture analyses (Pyron et al. 2020). At sympatric sites, we attempted to sample both black-bellied and shovel-nosed specimens. We also included one specimen each of the black-bellied *D. amphileucus* and *D. gvnigeusgwotli* from the Nantahala lineage as distantly related outgroups.

We extracted genomic DNA from tail-tip or liver tissue using a DNeasy Blood and Tissue Kit (Qiagen Inc.). We generated genomic libraries using the genotype-by-sequencing protocol (GBS; Elshire et al. 2011) at the University of Wisconsin-Madison Biotechnology Center with the ApeKI enzyme (New England Biolabs) from ∼50 ng of genomic DNA, sequenced on a NovaSeq6000 (Illumina Inc.). After demultiplexing, coverage ranged from 4–16 million reads per sample, with a mean of 9.4. The GBS assemblies were optimized in ipyrad v.0.7.30 (Eaton and Overcast 2020) under criteria proposed by Ilut et al. (2014) and McCartney-Melstad et al. (2019), yielding an optimal threshold of 96% similarity for clustering (Fig. S4). We allowed no barcode mismatches and used the “strict” adapter filtering option, leaving all other parameters as default values. The raw low-coverage assembly yielded 5,456 loci at a minimum of 50% locus coverage. After filtering for missing data (>20%) and singletons (Linck and Battey 2019), we retained 2,130 SNPs with 4,345 alleles for the ingroup assembly used in the clustering analyses. We dropped one *D. marmoratus* sample (RAP1949) based on low depth. All computational analyses were carried out on the GW HPC *Pegasus* cluster (MacLachlan et al. 2020). Accession numbers are given in Appendix S2 (Dryad repository https://doi.org/10.5061/dryad.rr4xgxdbp).

### Structure and Introgression

These analyses were performed only on the GBS data. We used a model-free geometric approach (DAPC; Jombart et al. 2010) for an initial estimate of population structure, which clusters individuals based on low-dimensional genetic distances calculated from PCA performed on SNP matrices. For DAPC clustering, selection of *K* using minimum BIC yielded an elbow curve with an optimal value of 4 (Fig. S6). We then estimated admixture and hybridization using two algorithms. First, we used sNMF (Frichot et al. 2014), which approximates the “admixture” model of STRUCTURE (Pritchard et al. 2000) and yields individual ancestry coefficients. We chose the value of *K* that minimized median cross-entropy across 100 replicates. We tested *K* values from 1–8, the total number of possible lineages. We then optimized the regularization parameter α for values 1, 5, 10, 50, 100, 500, and 1000, to minimize median cross-entropy across 100 replicates across all *K* values. This yielded values of α = 5 (Fig. S7) and *K* = 4 (Fig. S8).

We also estimated introgression across lineages represented as excess allele-sharing between branches using the Patterson’s *D* (ABBA-BABA) and *f_4_-*ratios to calculate the *f_b_*(C) statistics in the package ‘Dsuite’ (Malinsky et al. 2018, 2021). This tree-based approach estimates hybrid ancestry on a topology, inferred from the expected frequency distributions of site patterns under ILS versus reticulation, represented as the proportion of excess alleles shared between branch *b* and species C. This method is designed to estimate gene flow between populations but can be applied to closely related species as long as 1) species share a substantial amount of genetic variation due to common ancestry and ILS, 2) recurrent or backward mutations are negligible, and 3) substitution rates are uniform across species in the group. Dsuite integrates over all 4-taxon subtrees from a given phylogeny, allowing for the inference of multiple hybridization events, potentially including scenarios representing non-level-1 networks. Correspondingly, these estimates facilitate heuristic inference of the location and direction of hybrid edges placed in the estimated higher-level network. For the input topology, we used the backbone tree estimated from the AHE data in our network analyses (see above).

### Morphology

We measured 17 linear morphometric variables and a binary phenotype character for 335 specimens from 105 localities ranging from larvae to adults (16–85mm SVL), including all specimens in the GBS dataset (Appendix S2). These included all seven species-level lineages: *Desmognathus kanawha* (*n =* 63), *D. mavrokoilius* (*n =* 88), *D. intermedius* (*n =* 59), *D. marmoratus* (*n =* 46), *“quad”* C (*n =* 42), *“marm”* G (*n =* 12), and *“quad”* G (*n =* 25). All were assessed for the qualitative external morphological characteristics (Martof 1962) to code phenotype, with internal nares verified for most specimens. Thus, while overall phenotypes are discrete and binary based on apparently fixed diagnostic features (e.g., internal nares), specimens may still overlap in size and shape, and subtle differences within each phenotype may have evolved across lineages. We quantified phenotypes using 17 linear morphometric measurements adapted from Bingham et al. (2018) as described in our recent analyses (Pyron and Beamer 2022, 2023). We photographed the specimens on a copy stand at a 90-degree angle with a 60mm macro lens. We used the tps suite (Rohlf 2015) to place a series of 26 landmarks on the cranial and ventral surface of each specimen with a 20mm scale bar. Using the geomorph package in R (Adams and Otárola-Castillo 2013), we extracted 17 measurements in mm: SVL (snout-vent length), TL (tail length), AG (axilla-groin length), CW (chest width), FL (femur length [rather than hindlimb length]), HL (humerus length [rather than forelimb length]), SG (snout-gular length), TW (tail width at rear of vent), TO (length of third toe), FI (length of third finger), HW (head width), ED (eye diameter), IN (internarial distance), ES (eye-snout distance), ON (orbitonarial distance), IO (inter-orbital distance), and IC (inter-canthal distance).

We performed a simple set of linear morphometric analyses to assess divergence in phenotype across lineages. We first tested for significant differences in size and shape via MANCOVA with Pillai’s trace in R, using both genetic lineage and binary phenotype as response variables. We then re-projected the log-transformed measurements using principal components analysis and performed linear discriminant analysis to assess classification accuracy by clade. These analyses reflect qualitative ecomorphology and overall size, to quantify absolute morphometric differences. We augmented this with an allometric approach to characterize potentially subtle overlap in relative shape between lineages. We used the R package GroupStruct of Chan and Grismer (2021, 2022) for lineage-specific scaling of the 16 traits regressed against SVL using the 230 metamorphosed specimens >45mm SVL, the size of the largest larva. We then performed an additional allometrically-scaled LDA on SVL + 16 size-corrected traits by lineage, excluding the qualitative phenotypes. While some degree of size and shape variation is known across these populations already (Martof 1962; Pyron and Beamer 2022), we hypothesize that the two distinct phenotypes will be completely segregated morphometrically. Alternatively, if intermediate, novel, or transgressive phenotypes reflect the hybrid origin of some lineages, we hypothesize that these populations will overlap their parental species morphometrically, with some specimens being mis-classified by the LDA as one or more of the ancestral groups contributing to the originating hybrid event.

## Results

### Network Estimation

We analyzed the AHE data representing 233 gene trees and 122 individuals from Pyron et al. (2022) following the steps outlined in the Methods and the SI. Table S3 shows the quartet trees associated with those networks identified as T-quartets, while Table S4 shows them for the networks identified as B-quartets. Following the procedure outlined in the SI, we repeated our approach exhaustively on all seven-taxon unrooted trees, resulting in the two trees depicted in Figure S12. One of such is the tree depicted in gray in Figure 4, which we propose as the “backbone” of the species network because it is identical to the previous concatenated and species-tree analyses of different AHE datasets (Pyron et al. 2020, 2022). We are therefore confident in this arrangement of the seven lineages as representing the most accurate bifurcating consensus topology. The choice of such a tree should not affect the species network inference given both trees differ only on the placement of *“quad”* C and are only a Nearest Neighbor Interchange move (NNI) from each other and both are displayed in the resulting network. By combining the trees obtained from our method, the signals of quartets identified as B-quartets, and the hybrid signals obtained from sNMF and Dsuite, we then constructed a network (Fig. 5).

As mentioned above, the selected backbone tree was used as the input for Dsuite, which most strongly supports a hybrid event between *Desmognathus mavrokoilius* and *D. marmoratus,* also supported by the sNMF analysis of the GBS data (Fig. 3) and our previous Dsuite analysis of the AHE data (Pyron et al. 2022). Given the structure of the backbone tree, this implies that there is a hybrid edge between the terminal edges of *D. mavrokoilius* and *D. marmoratus*. This is also corroborated by mitochondrial exchange (Kozak et al. 2005). Based on the information from the B-quartets, we propose that the direction of such a hybrid edge must be towards *D. marmoratus*, implying that it is a hybrid descendant. Additional evidence (see below) also favors *D. marmoratus* arising via hybridization. This hybridization event is depicted in Figure 4, where the backbone tree is shaded in gray and the edge added for this event is labeled *a*.

**Figure 3.**
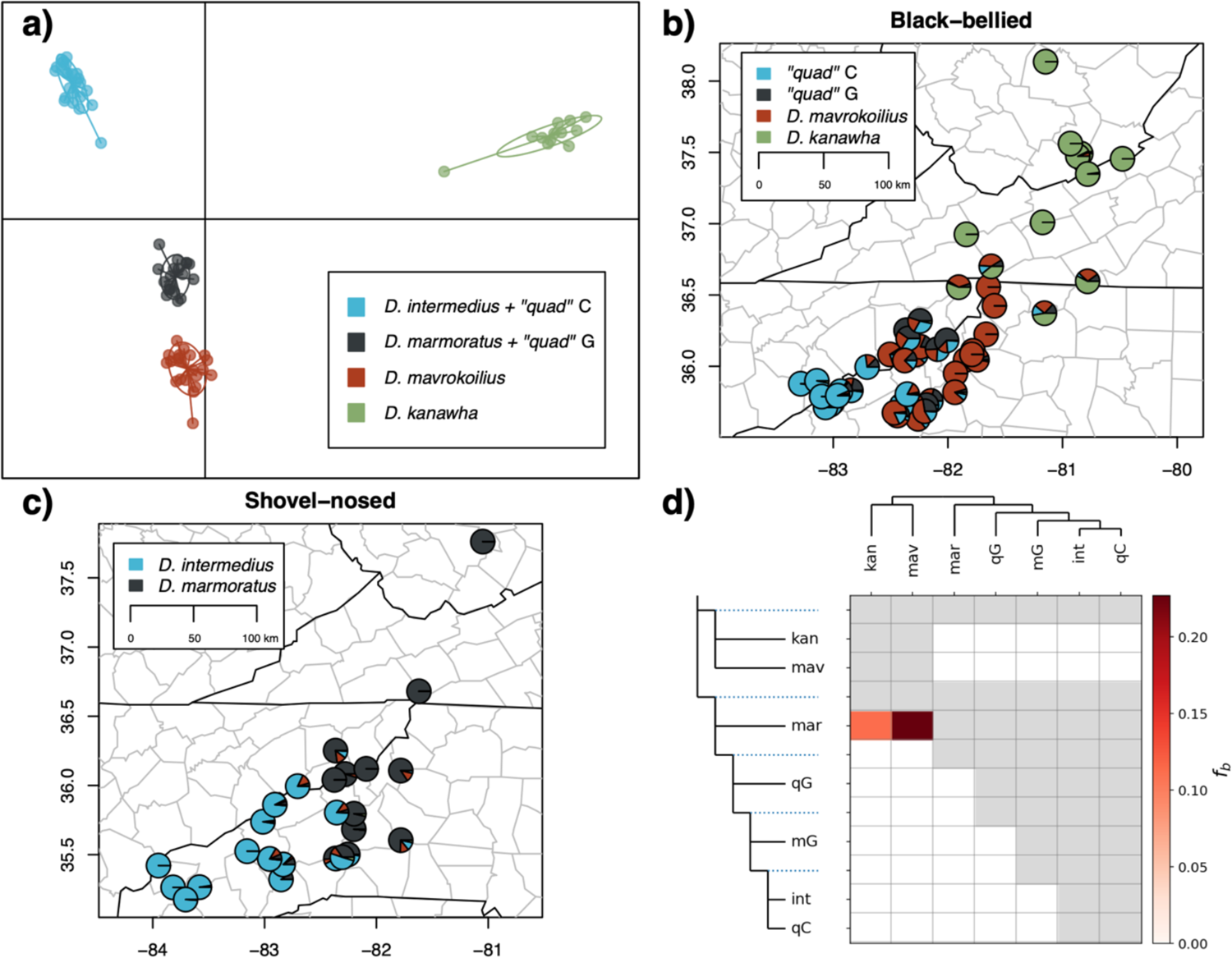
Results from DAPC (a), sNMF (b, c), and Dsuite (d) analyses of the SNP matrix from the GBS sequencing run, showing spatial and phylogenetic patterns of genetic clustering and admixture. The localities are again divided by phenotype (see Fig. 2), showing that populations genetically clustering with *Desmognathus intermedius* and *D. marmoratus* (including *“marm”* G) exhibit both shovel-nosed and black-bellied (*“quad”* C and G) morphologies at some localities. Dsuite identifies substantial introgression into *D. marmoratus* from both *D. kanawha* and *D. mavrokoilius*. In (b) and (c) the pie charts represent individual ancestry coefficients. In (d), the matrix entries represent the proportion of excess alleles (*f_b_* statistics) shared between the donor (column) and recipient (row) branches across the phylogeny.

Both the admixture signal from sNMF using the AHE data (Pyron et al. 2022) and the sNMF analysis of the GBS data (Fig. 3) suggest that there is a hybrid event between *Desmognathus marmoratus* and *“quad”* G. This hybrid event is depicted with the edge labeled *b* in Figure 4. If such an edge is directed as depicted, the resulting network agrees with two rows (19,20) in Table S4. As we discuss below, there is also evidence suggesting that *“quad”* G is the descendant of the hybridization event. The sNMF analysis of the GBS data (Fig. 3) also shows that *“quad”* G contains ancestry from *D. mavrokoilius* (also supported by Dsuite and sNMF using the AHE data; Pyron et al. 2022), and the group that represents *D. intermedius* + *“quad”* C. This is the scenario depicted by edges *a* and *b* in Figure 4, where the signal of *D. mavrokoilius* comes from the earlier hybridization with *D. marmoratus*. Moreover, when combining edges *a* and *b,* the network is in agreement with three more rows (16,26,27) in Table S4.

**Figure 4.**
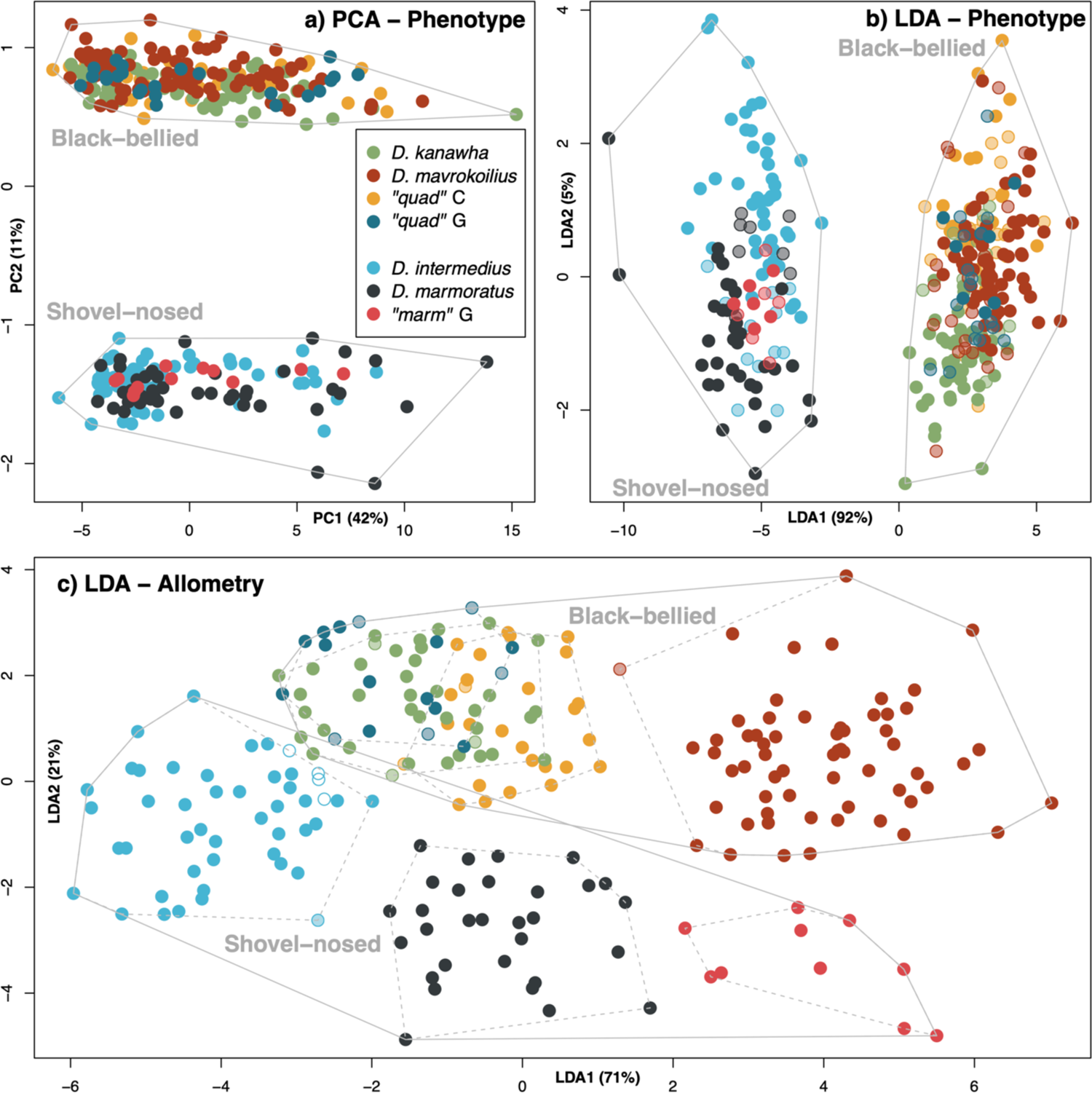
Results from PCA (a) of all phenotypic and morphometric variables, LDA (b) of the 17 linear measurements showing strong differentiation between phenotypes but limited differentiation between lineages, and (c) LDA based on allometric size-correction of shape across lineages without discrete phenotypes. In the LDA plots, solid circles were correctly classified to genetic lineage, semi-transparent were mis-classified between lineages within phenotypes, and open circles were mis-classified between phenotypes. Convex hulls are drawn solid around the two phenotypes and dotted around each genetic lineage in (c). Allometric scaling (c) shows a small overlap in the total morphospace between the two phenotypes, but not between any of the individual black-bellied and shovel-nosed genetic lineages, which are completely differentiated along the first two axes except for *D. kanawha* and *“quad”* C & G.

**Figure 5.**
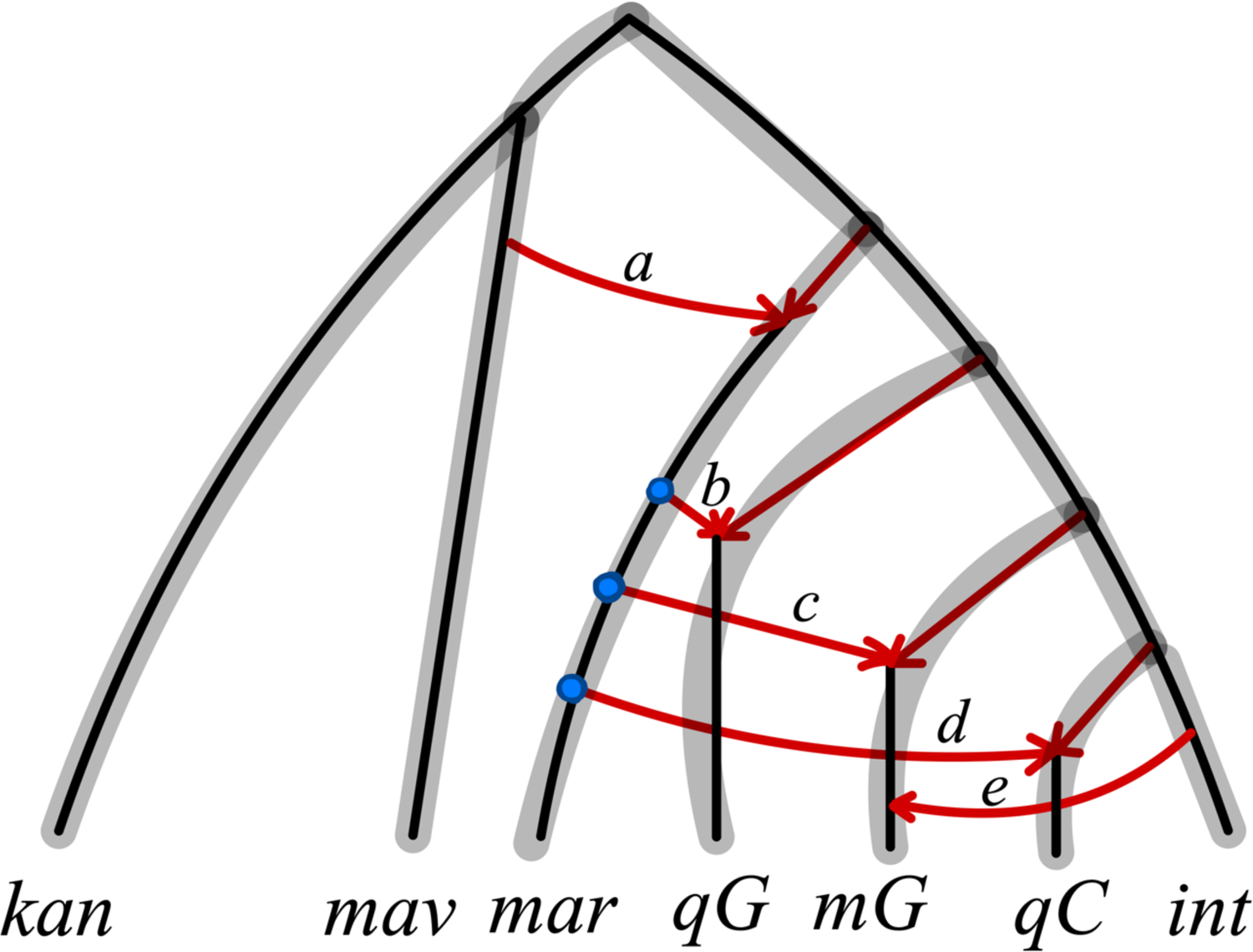
Phylogeny (gray) and network (red) estimated from congruent signals independently in the AHE and GBS datasets using our new heuristic method. Based on known patterns of mitochondrial exchange and the sNMF and Dsuite analyses detailed above, combined with the signals from B-quartet networks in Table S4, we propose that the species network contains the hybrid events depicted in red. This network agrees with the largest proportion of gene-tree signals across all analyses. The order of the blue nodes in the edge with terminal taxon *Desmognathus marmoratus* are chosen according to the order of their corresponding hybrid event in the edge with taxon *D. intermedius*. The blue nodes may not have appeared in this exact sequence, but our method cannot determine the precise ordering. Abbreviations at tip labels are: *kan – D. kanawha, mav – D. mavrokoilius, mar – D. marmoratus, qG – “quad”* G, *mG – marmoratus* G, *qC – “quad”* C, and *int – D. intermedius*.

From the same admixture analyses, we obtain two other hybrid events. One suggests that *“marm”* G is a mixture of *Desmognathus intermedius* and *D. marmoratus*. This can be depicted by adding a hybrid edge to the backbone from *D. marmoratus* to *“marm”* G as depicted in Figure 4, using the edge labeled *c*. For this hybrid event, there is no ambiguity on the direction given that sNMF already displays the parents as such: *D. marmoratus* is the pre-existing parental lineage, and the admixture analyses recover *“marm”* G as a hybrid population between *D. intermedius* and *D. marmoratus*. This event agrees with one row (12), and when combined with the other events already mentioned, agrees with 3 more rows (13,23,25), as seen in Table S4.

The final hybridization event obtained from sNMF analysis of the GBS data suggests that *“quad”* C is a mixture of *Desmognathus intermedius* and small amounts (<23%) of estimated genomic ancestry from *D. marmoratus* and *D. mavrokoilius*. We propose that there is a hybrid edge from *D. marmoratus* to *“quad”* C in the backbone tree. This is depicted by the edge labeled *d* in Fig. 5. These edges agree with 5 more rows (1,9,10,17,18) in Table S4. We therefore hypothesize that the signal of *D. mavrokoilius* could come from the previously mentioned hybridization between *D. mavrokoilius* and *D. marmoratus*, which would also explain the phenotypic transplant into that lineage. When combining this hybridization and the other described above such a network has four more rows (11,14,15,24) in agreement with Table S4.

Finally, we add the hybridization from *D. intermedius* to *“marm”* G, labeled *e* in Fig. 5, which is obtained from the other tree obtained from our analysis (Fig. S12 right). The resulting network harmonizes the signals obtained from Dsuite and sNMF and displays 53 out of the 62 quartet trees (85%) associated with all quartet networks (Tables S3 and S5). There are 9 quartet trees that are not in agreement with the network. This implies that either hybrid events are missing, or that these events were resolved incorrectly. In any case, since there is no more supporting signal from Dsuite or sNMF, we refrain from interpreting them further. In terms of the final network (Fig. 5), we tried different configurations of hybridizations, but none were able to obtain the same level of agreement with the B-quartets, Dsuite, and sNMF analyses. Given the complex nature of the space of networks, this cannot reasonably be done exhaustively over all possible combinations. Finally, we cannot exclude the possibility that there are still other hybrid events missing from the network (both internal and external), but for now we have no way of recovering them independently.

### Structure and Introgression

A PCA plot (Dray and Dufour 2007) reveals partial to complete separation of the 7 mito-nuclear candidate species (Pyron et al. 2020, 2022), with overlap between *D. intermedius* & *“quad”* C, and between *D. marmoratus* & *“quad”* G (Fig. S5). The DAPC analysis yielded strong support for *K=*4 (Fig. 3) with a distinct elbow curve using BIC (Fig. S6). These four clusters consist of *Desmognathus kanawha* (part), *D. mavrokoilius* (plus two hybrids with *D. kanawha*), *D. marmoratus* + *“marm”* G + *“quad”* G, and *D. intermedius* + *“quad”* C. These four groups form non-overlapping clusters on the discriminant axes, and all samples have membership probabilities ∼1. Taken literally, this would imply the existence of four hybridizing parental species, two of which are phenotypically dimorphic, but which display little morphological intermediacy between these phenotypes (see below). While placement of *“marm”* G was uncertain in previous analyses (Pyron et al. 2022), it is recovered firmly as a population of *D. marmoratus* here, with only some genomic ancestry from *D. intermedius*.

The sNMF optimization yielded essentially the same clusters as the DAPC analysis. While being cautious not to over-interpret this potentially over-simplified model (Lawson et al. 2018), the estimation of individual ancestry coefficients lends a modestly more coherent explanation to the phenotypic and topological patterns revealed by previous analyses. The geographic context of the individual ancestry coefficients yields a relatively straightforward interpretation. Clines of genomic admixture occur along the Appalachian Mountains in a southwest to northeast direction between parapatric sister lineages (*Desmognathus intermedius/D. marmoratus* and *D. kanawha/D. mavrokoilius*). Similarly, introgression is observed in sympatry at several localities between two non-sister lineages (i.e., *D. marmoratus* and *D. mavrokoilius*). In the Dsuite analysis, the matrix of *f_b_* values can be read to indicate the proportion of excess alleles shared between a donor species in the columns and the recipient branch in the rows (Malinsky et al. 2021). We estimate one recipient branch involved in gene flow from *D. mavrokoilius* to *D. marmoratus*. Patterns from all three analyses are consistent with previous mitochondrial evidence of exchange between *D. intermedius* and *“quad”* C, and between *D. marmoratus* and *D. mavrokoilius* (Kozak et al. 2005; Beamer and Lamb 2020).

### Morphology

The PCA of all morphometric and phenotypic variables reveals strong separation between phenotypes and limited differentiation within phenotypes between genetic lineages (Fig. 4a, b). The MANOVA indicated significant differences in external morphometrics when comparing both phenotype (*F*_1,333_ = 29.5, Pillai = 0.61, *P* = <0.000001), and genetic lineage (*F*_6,328_ = 5.3, Pillai = 1.3, *P* = <0.000001) using the 17 variables. The LDA classifier was 100% accurate by phenotype, though this is trivial given that the model included potentially diagnostic characters. As expected, total accuracy is lower by genetic lineage at 69%; this is driven by misidentification within shovel-nosed (10–16%) and black-bellied (19–32%) lineages. Thus, strong separation between the two phenotypes is still observed when classifying by genetic lineage, but much less discrimination is possible between lineages of each phenotype (Fig. 4). Other putatively diagnostic traits (e.g., tail height) were not measured and would likely improve accuracy. Using allometric scaling to correct for relative size increases discrimination between lineages (Fig. 4c), while still supporting the underlying binary phenotypic dimorphism. The classifier is 93% accurate overall, with only 0–2% misclassification for shovel-nosed lineages, and 0–12% for black-bellied. No black-bellied specimens are classified as shovel-nosed, while four *D. intermedius* are classified as *D. kanawha* (3 specimens) or *“quad”* G (1 specimen).

There is no overlap or misclassification between *D. intermedius* and *“quad”* C, or *D. marmoratus* and *“quad”* G. The three shovel-nosed lineages occupy a unique region of morphospace, while *D. mavrokoilius* occupies a distinct area from the overlapping *D. kanawha, “quad”* C, and *“quad”* G. Consequently, we reject the hypothesis that hybrid lineages exhibit intermediate or overlapping phenotypes between their parentals (or novel or transgressive morphologies), in both absolute and relative terms. Instead, these results favor a model where black-bellied and shovel-nosed populations are phenotypically differentiated in two distinct regions of morphospace, within which each lineage may exhibit subtle divergence.

## Discussion

### Higher-Level Networks

Inferring evolutionary relationships in the presence of hybridization presents many challenges (Huson and Bryant 2006; Huson et al. 2010; Morrison 2011; Folk et al. 2018), and gene-tree incongruence driven by incomplete lineage sorting (ILS) can be difficult to distinguish from that driven by hybridization (Nakhleh 2010; Olave et al. 2018; Hahn and Hibbins 2019).

Much work has been done to reconstruct species networks, particularly those that are level-1. SNaQ (Solís-Lemus and Ané 2016) and NANUQ (Allman et al. 2019) made great advancements in estimating level-1 networks in the presence of ILS. However, statistically identifiable methods are limited exclusively to level-1 networks. While obtaining identifiability for level-1 networks was a substantial achievement, more datasets like the one presented here are arising which violate level-1 assumptions. Unfortunately, as pointed out by Pardi and Scornavacca (2015) multiple higher-level networks may be represented by the same set of trees, suggesting that in some cases we may lack the ability to resolve alternative hypotheses. How to generalize methods for level-1 resolution to a broader family of higher-level networks has been unclear. In this work, we present a novel heuristic approach to detecting meaningful signals for overlapping hybrid edges from higher-level networks (displayed trees) from gene trees and other data.

We address this challenge by differentiating between T-quartet and B-quartet networks (networks that display only one tree topology or display multiple tree topologies, respectively) based on a given set of gene trees under the NMSC. These quartets are then used to heuristically obtain displayed trees on networks. Extrinsic data from sources such as mitochondrial exchange, individual ancestry coefficients, and excess allele-sharing under the MSC may all be useful in breaking ties and selecting networks with the greatest congruent signal in terms of explaining the greatest number of B-quartets as detailed above. Our approach offers a potential advance by recovering features from higher-level networks, which in some cases are sufficient to uniquely identify the full network. As with most methods, ours is impacted by gene-tree error. Another source of error is the violation of the assumption that the trees associated with the largest CFs are displayed in the networks. While this should be true in most cases, it can be false. Solís-Lemus and Ané (2016) showed that under the NMSC, there are level-1 networks where the most dominant gene tree quartet is not displayed in the network. Nonetheless, Allman et al. (2019) argued that the set of parameters required for this to occur (extremely short branches or very large populations) must be extreme, and unlikely to be common in most empirical scenarios.

We conducted several small simulation studies of different networks with various parameters (edge lengths and hybridization probabilities; see SI). Initial simulation results with 5–7 terminals and 1–2 B-quartets suggest that this method is consistent in cases with approximately this level of complexity (Fig. 1; Appendix S1). In all simulations, our method recovered unrooted displayed trees, that is, unrooted trees in agreement with the network. In some cases, these signals were enough to recover the full network topology. As with all existing coalescent-based methods, our method cannot identify small cycles (those of sizes 2 and 3) but performs well on cycles greater than 4. Future work could extend this approach to develop a statistically consistent method to infer higher-level networks in the presence of ILS using information from B-quartets and T-quartets. An important first step would be to determine what features are theoretically identifiable under these conditions.

### Cryptic Speciation and Phenotypic Diversity

Clustering analyses estimate four major parental lineages comprising both black-bellied (*Desmognathus kanawha* and *D. mavrokoilius*) and shovel-nosed (*D. intermedius* and *D. marmoratus*) species. Within these reside three “minor” populations (*“marm”* G and *“quad”* G nested within *D. marmoratus* and *“quad”* C nested within *D. intermedius*) of at least partial hybrid origin. They are differentiated from their parentals by distinct mitochondrial haplotypes (Beamer and Lamb 2020) and topological positions in phylogenomic analyses (Pyron et al. 2020, 2022) as well as geography (*“marm”* G) and phenotype (*“quad”* C & G). Based on their evident degree of genetic, geographic, and ecomorphological distinctiveness, we conclude that they are relatively “ancient” (i.e., >1–2Ma; Kozak et al. 2005) and well-established or stabilized lineages, rather than fleeting or ephemeral populations representing instances of secondary contact resulting in recent admixture or introgression.

We also find the apparent presence of admixed genomes suggesting hybrid ancestry between both sister (*D. kanawha/D. mavrokoilius* and *D. intermedius/D. marmoratus*) and non-sister (*D. mavrokoilius* and *D. marmoratus*) lineages. This indicates a complex history of reticulation requiring a non-level-1 evolutionary network to visualize, in which hybrid edges were either shared or overlapping across multiple introgression events (Fig. 5). These reticulations occur both between and among phenotypically distinct black-bellied and shovel-nosed lineages. The basis of this phenotypic dimorphism is unclear; we suspect that it is a polygenic threshold trait since it is binary and manifests across multiple skeletal and soft tissue features, but it is not associated with a clear pattern of genomic ancestry in this or previous analyses (Beamer and Lamb 2020; Pyron et al. 2020, 2022).

In salamanders such as *Plethodon*, some intraspecific phenotypic polymorphisms seem linked to one or a small number of loci and are associated with drift-based fixation in or around glacial refugia, without obvious ecomorphological selection (Thurow 1961). In *Desmognathus,* some phenotypes appear to stem from frequency dependent Batesian mimicry (Labanick 1983). In contrast, polygenic threshold traits such as paedomorphosis in *Ambystoma* have a complex genomic basis (Roff 2008). As paedomorphic *Ambystoma* populations are obligately aquatic (contrasted with terrestrial or semi-aquatic metamorphs), previous workers hypothesized that the repeated evolution of paedomorphosis may have a strong effect in promoting speciation (Percino-Daniel et al. 2016). However, recent range-wide genomic analyses suggested that geography exerts a stronger effect on lineage formation than phenotype (Everson et al. 2021). In contrast here, hybridizing parental species with different phenotypes (the ancestral black-bellied and shovel-nosed lineages) produce both allopatric (*“marm”* G) and parapatric (*“quad”* C and *“quad”* G) daughter lineages, which subsequently backcross with their parentals.

Consequently, we allocate *“marm”* G to *D. marmoratus* with which it shares the majority of its genomic ancestry based on the GBS data. We also retain *“quad”* C and G within *D. mavrokoilius* as previously suggested by Pyron and Beamer (2022). The latter action is nomenclaturally conservative but does not fully resolve the taxonomic status of these populations. While they do not have a unique genetic composition in our GBS dataset, they occupy topologically distinct positions in mitochondrial and AHE analyses (Beamer and Lamb 2020; Pyron et al. 2020). In contrast, they also exhibit admixture with *D. intermedius, D. marmoratus,* and *D. mavrokoilius* (see Results; Pyron et al. 2022) in parapatry or sympatry with those species (Fig. 2, 3). Furthermore, they overlap morphometrically not with *D. mavrokoilius,* but with *D. kanawha* (Fig. 4), though the distinct phenotypes of some *“quad”* C populations are thought to simply reflect local adaptations to higher elevations (Beachy and Bruce 2003).

Resolving the systematic treatment of such populations decisively can be difficult or impossible; they fulfill some, but not all, criteria for hybrid speciation (Gross and Rieseberg 2005; Mallet 2007; Schumer et al. 2014). They clearly maintain unique and independent evolutionary identities, genetically or morphologically differentiated from at least one parental lineage in sympatry or parapatry. Yet, they also exhibit genetic interactions with one or more parental lineages where they are in either primary (implying incomplete divergence) or secondary (implying backcrossing) contact. Finally, genomic differentiation is evident in some markers (e.g., mitochondria, AHE) but not others (e.g., GBS), and they appear to occupy identical adaptive zones to at least one parental lineage as indicated by phenotype. While such patterns have not yet been widely reported in the literature, similar instances in eels and lampreys have been attributed to incomplete reproductive isolation (Rougemont et al. 2015; Barth et al. 2020), which may be an adaptive optimum in some instances (Servedio and Hermisson 2020) and even promote diversification (Seehausen 2004; Svardal et al. 2020).

### Ecomorphological Hybrid Speciation

The pattern observed in these salamanders appears to correspond with recent widely accepted definitions of homoploid hybrid speciation (Mallet 2007; Abbott et al. 2010). For instance: “(1) reproductive isolation of hybrid lineages from the parental species, (2) evidence of hybridization in the genome, and (3) evidence that this reproductive isolation is a consequence of hybridization” (Schumer et al. 2014). More permissive definitions have been proposed (e.g., Nieto-Feliner et al. 2017), but we suggest that even under the strict interpretation of Schumer et al. (2014, 2018), the Pisgah clade satisfies their three criteria. We observe (i) at least partial genetic isolation between sympatric hybrid and parental lineages separated by phenotypically mediated microhabitat selection, (ii) multiple genomic signatures of hybridization across the nuclear and organellar genomes, and (iii) evidence that the phenotypic shift in hybrid lineages is the key factor facilitating their genetic isolation from parental backcrossing. However, due to the unusually conserved nature of the dimorphic ecological morphotype, the Pisgah clade differs from other well-known empirical instances of hybrid speciation in several distinct ways.

Several of the named lineages including *Desmognathus intermedius, D. marmoratus, “marm”* G and *“quad”* C & G appear to have arisen entirely or in part via hybrid genomic contributions from genetically distinct ancestral lineages. In at least the case of *“quad”* C & G, subsequent parental and hybrid distinctiveness was maintained by a change in phenotype causing ecological niche partitioning. Consequently, diversification in the Pisgah clade appears to reflect ecomorphological hybrid speciation through the transmission of a polygenic threshold trait via repeated reticulation events between parental lineages in a hybrid swarm (Seehausen 2004). In contrast to classic systems such as sticklebacks (Hatfield and Schluter 1999) and butterflies (Duenez-Guzman et al. 2009), the Pisgah ecomorphs of *Desmognathus* are sympatric or syntopic, but exhibit replicated bimodality in strongly differentiated habitat (stream bed versus streamside) and resource (e.g., diet; Martof and Scott 1957; Davic 1991) usage.

Consequently, there is little or no morphological intermediacy; instead, we observe a discrete phenotypic dimorphism across multiple lineages. Hybrid fitness is demonstrated by the common observation of admixed genotypes between phenotypes; morphological shovel-nosed (aquatic) with substantial black-bellied (semi-aquatic) ancestry and vice versa. We propose that the initiation of this mechanism occurs via transmission of threshold alleles at some number of loci resulting in a phenotypic switch between shovel-nosed or black-bellied morphologies in the resulting hybrid population. Subsequent ecological selection then maintains ecomorphological distinctiveness of the dimorphic phenotypes in non-overlapping microhabitats and resource niches. The high observed frequency of admixed genotypes also suggests limited post-zygotic isolation and an unknown impact of reinforcement via sexual incompatibility or mate choice (Arnold et al. 1993); nothing is known of sexual behavior in these species.

Existing theories of ecological hybrid speciation (Gross and Rieseberg 2005) typically posit hybrids that are either intermediate, novel, or transgressive in comparison to parentals in their phenotype, ecology, and fitness (Seehausen 2004; Hendry et al. 2007; Marques et al. 2019). Here, we find that hybrids are essentially isomorphic with at least one parental lineage (Short and Streisfeld 2023); hybrids are moving back and forth between existing binary fitness peaks. We also see a curious asymmetry in transmission of threshold alleles for the phenotype switch (Fig. 5). Given that the basal split in the tree is between black-bellied (leading to *Desmognathus kanawha + D. mavrokoilius*) and shovel-nosed (leading to *D. intermedius* + *D. marmoratus*) phenotypes, hybridization between *D. mavrokoilius* and the ancestral shovel-nosed yield *D. marmoratus*, but subsequent hybridizations involving the *D. marmoratus* branch yield both shovel-nosed (*“marm”* G) and black-bellied (*“quad”* C and *“quad”* G) phenotypes. This suggests that the phenotype is linked to a small number of loci, probably in epistasis, since a genomically distinct source of black-bellied ancestry is lacking in *“quad”* C and *“quad”* G.

In some Lake Malawi cichlids, hybridization produces phenotypic dimorphism depending on the heterozygosity of two epistatic loci (Parnell and Streelman 2013). However, those traits are linked to sex determination and sexual signaling by phenotype, distinct from the scenario here where there is no apparent sexual dimorphism. Contrastingly, hybridization between closely related sculpin species produced a novel ecomorphotype yielding adaptive potential for habitats previously unoccupied by either parental species (Nolte et al. 2005). In this *Desmognathus* system, phenotypic dimorphism apparently results in the reinvasion of one of two distinct habitats: stream bed versus streamside. This allows ecologically differentiated populations to exist in close geographic proximity (oftentimes even in the same streams) with apparently reduced or limited gene flow. Subtle variation within each phenotype (e.g., dwarfism in *“quad”* C) seems to result from subsequent local adaptation (Beachy and Bruce 2003).

A final major difference between many previous systems and our study is that the phenotypes themselves are relatively ancient and seemingly unaltered by repeated hybridization. In fact, the historical origin of the two phenotypes themselves remains unresolved; both the shovel-nosed and black-bellied phenotypes are shared with four species (three black-bellied and one shovel-nosed) in the distantly related and earlier diverging Nantahala clade (Pyron et al. 2020). Mitochondrial data support monophyly of the Nantahala + Pisgah lineages (Kozak et al. 2005; Beamer and Lamb 2020), indicating a deep-time reticulation prior to the diversification of the extant lineages (∼8–15 Ma; Kozak et al. 2005) which has yet to be detected using tree- or SNP-based methods (see Pyron et al. 2020). Some similar instances have recently been demonstrated in alpine lake fishes (De-Kayne et al. 2022) and monkeyflowers (Short and Streisfeld 2023), where selection has removed much of the signal of ancient introgression that resulted in phenotypic shifts between hybrid lineages through time. Identifying the genetic basis of these black-bellied and shovel-nosed phenotypes, potential subtle differences between the genetic lineages of each, and their phylogenetic origin will be crucial to untangle the complex evolutionary history of the group. Consequently, a more complete genomic perspective on trait origins and ancient introgression may provide substantial insight into ecomorphologically mediated mechanisms of hybrid speciation in these salamanders.

## Acknowledgements

The authors used the University of Wisconsin – Madison Biotechnology Center’s DNA Sequencing Facility (Research Resource Identifier – RRID:SCR_017759) to generate GBS libraries and sequence GBS libraries. This work was completed in part with resources provided by the High Performance Computing Cluster at The George Washington University (Information Technology, Research Technology Services), and the Smithsonian High Performance Cluster (SI/HPC), Smithsonian Institution (https://doi.org/10.25572/SIHPC). We thank I. Sanmartín, C. Solís-Lemus, and four anonymous reviewers for their comments.

## Data Availability

The underlying data are available in Dryad Repository https://doi.org/10.5061/dryad.rr4xgxdbp, and the Supplemental Information and code for the network estimation approach can be found at Zenodo repository https://doi.org/10.5281/zenodo.10386850 and https://github.com/kyleaoconnell22/high-level-networks-desmog.

